# DNA Double Strand Breaks cause chromosome loss through sister chromatid tethering in human embryos

**DOI:** 10.1101/2022.03.10.483502

**Authors:** Jenna Turocy, Diego Marin, Shuangyi Xu, Jia Xu, Alex Robles, Nathan Treff, Dieter Egli

## Abstract

Genome editing by DNA double-strand breaks (DSB) is currently being investigated as a tool to treat or even prevent heritable diseases^1^. However, DNA repair mechanisms in the human embryo remain poorly understood and DSBs may result in chromosome loss ^2,3^. Here we provide evidence of whole and segmental chromosome loss in over one third of chromosomes 16, 17 and X targeted by CRISPR/Cas9-induced DNA DSB, including pericentromeric and mid-arm sites. Chromosomal changes were asymmetric relative to the Cas9 cut site: segmental losses occurred on both centric as well as acentric chromosome arms, while gains were exclusively found on acentric arms, suggesting that centromeres in broken chromosomes continued to mediate sister chromatid separation. Using this pattern of chromosomal errors, we were able to define new genomic coordinates of the active centromere on chromosome 16. Asymmetry was also found in the attrition of gDNA at the break site: attrition occurred centromeric of the DSB, while telomeric to the break, chromosomal ends were protected. Thus, spindle forces at centromeres and end tethering and protection at DSBs are antagonistic forces that interfere with accurate segregation of sister chromatids. Thereby, a single DSB is sufficient to result in the loss of a chromosome from the embryo. These results highlight the risks of aneuploidy in CRISPR/Cas9 genome editing, while also providing a mechanism for mitotically acquired aneuploidy caused by DNA breaks in human embryos.

---

Studies since the early 1980s have demonstrated the power of using DNA double-strand breaks (DSB) for targeted genetic change, first in yeast and then in mammalian cells ^4,5^. In the 2000s, scientists began to discover how bacteria possess unique abilities to fight viruses by cutting the viral genes with a highly specific RNA guided nuclease ^6^. Researchers quickly adapted this technology known as “Clustered Regularly Interspaced Short Palindromic Repeats” or CRISPR to other species and research has continued to advance rapidly. Clinical trials are currently underway using postnatal somatic cells to modify a person’s DNA to treat or cure a disease ^7^. Somatic gene therapies however may be limited in their ability to reverse damage that has already occurred and reach the billions of cells that are needed to adequately treat the disease. Germline human genome editing, on the other hand, alters the genome of a human embryo at its earliest stages and may prevent disease permanently including in future generations.

For the therapy to be effective and safe, the precision of DNA recognition, cleavage, and repair must be extremely high, with exact in-frame changes and absence of off-target effects. DNA repair mechanisms, however, are error prone. CRISPR/Cas9 mediated studies in human embryos have shown that indels (small insertions or deletions) occur at a high frequency (reviewed in Turocy et al.^8^. In addition to indels, CRISPR/Cas9-induced DSB can result in more extensive genetic changes including chromosome loss. In a prior study using a guide RNA targeting a mutation in the *EYS* gene on chromosome 6, DNA breaks in approximately half of the embryos injected resulted in partial or whole chromosome loss ^2^. An off-target site on the arm of chromosome 16 similarly resulted in segmental chromosome losses ^2^. Fogarty and colleagues have also reported chromosome change at the *POU5F1* locus after CRISPR/Cas9 injection ^3^. Whether failure to reseal the DSB, or abnormal repair results in chromosomal change in human embryos is not currently known. Not all CRISPR/Cas9 mediated human embryo studies, however, have examined karyotypes and reported chromosome loss after CRISPR/Cas9 injection (reviewed in Turocy et al. 2021)^8^. Other repair outcomes, including interhomolog repair through gene conversion resulting in loss of heterozygosity have been suggested^9^ and have been demonstrated in mice^10^. It is unclear whether repair pathway choice and failure to repair the break is dependent on the chromosomal location. Given the potential catastrophic consequences of segmental or whole chromosome loss on the developing human embryo, further studies to understand DNA repair pathway choice and the frequency of chromosomal events after CRISPR/Cas9 injection are needed.

Prior work studying meiotic origins of human aneuploidy have demonstrated that DNA DSBs within specific regions of the genome are at increased risk of genome instability and chromosome missegregation. These vulnerable regions include telomeres, centromeres and peri-centromeres ^11–15^. It has also been observed that pericentromeric regions are more frequently involved in chromosome rearrangements and breaks in cancer cells (reviewed in Barra et al ^16^). Furthermore, microdeletions and microduplications near centromeres can give rise to disorders such as 16p11.2 microdeletion or 16p11.2 duplication syndrome which have been associated with an increased risk of intellectual disability and psychiatric disorders ^17^.

We hypothesized that a CRISPR-induced DNA DSB near the centromere of chromosome 16 or X would result in whole and segmental chromosome loss. These two chromosomes were chosen because they show frequent aneuploidies in human oocytes ^18^, providing a potential target for ploidy correction in the embryo and an *in vitro* model for spontaneous chromosome loss. To test this hypothesis, CRISPR/Cas9-guide RNAs (gRNA) were designed to target unique genomic sites in the pericentromeric region of Chromosome 16 and Chromosome X. Embryos were then analyzed on a single-cell level at the cleavage stage, and found to have whole and/or segmental chromosome loss in over one third of chromosomes targeted by CRISPR/Cas9. Human embryos injected with CRISPR/Cas9 gRNA targeting the mid-arm of chromosome X also demonstrated segmental chromosome loss exemplifying the risks of CRISPR/Cas9 induced DNA DSB regardless of gene location. The pattern of Cas9-induced chromosomal losses and gains allowed us to determine the genomic coordinates of the active centromere of chromosome 16. Chromosomal gains are found for acentric chromosomal arms, while both acentric and centromere containing chromosomal arms give rise to losses. Furthermore, one embryo with a spontaneous trisomy 16 due to gain in meiosis also showed chromosome 16 loss after CRISPR/Cas9 injection, which points to the possibility of targeted chromosome loss and future therapeutic applications.

## RESULTS

### DNA DSBs frequently result in chromosomal losses

CRISPR/Cas9-single-guide RNAs (sgRNA) were designed to target unique genomic sites in the pericentromeric region of Chromosome 16 and Chromosome X (**Fig 1a, Table S1)**. Thirty IVF-generated zygote embryos previously frozen at the 2PN stage were thawed and injected with Cas9 ribonucleoprotein (RNP) and guide RNA (gRNA) into the cytoplasm immediately after thawing **(Fig 1b)**. Survival rate post injection was 96.7% (29/30).

**Fig. 1.**
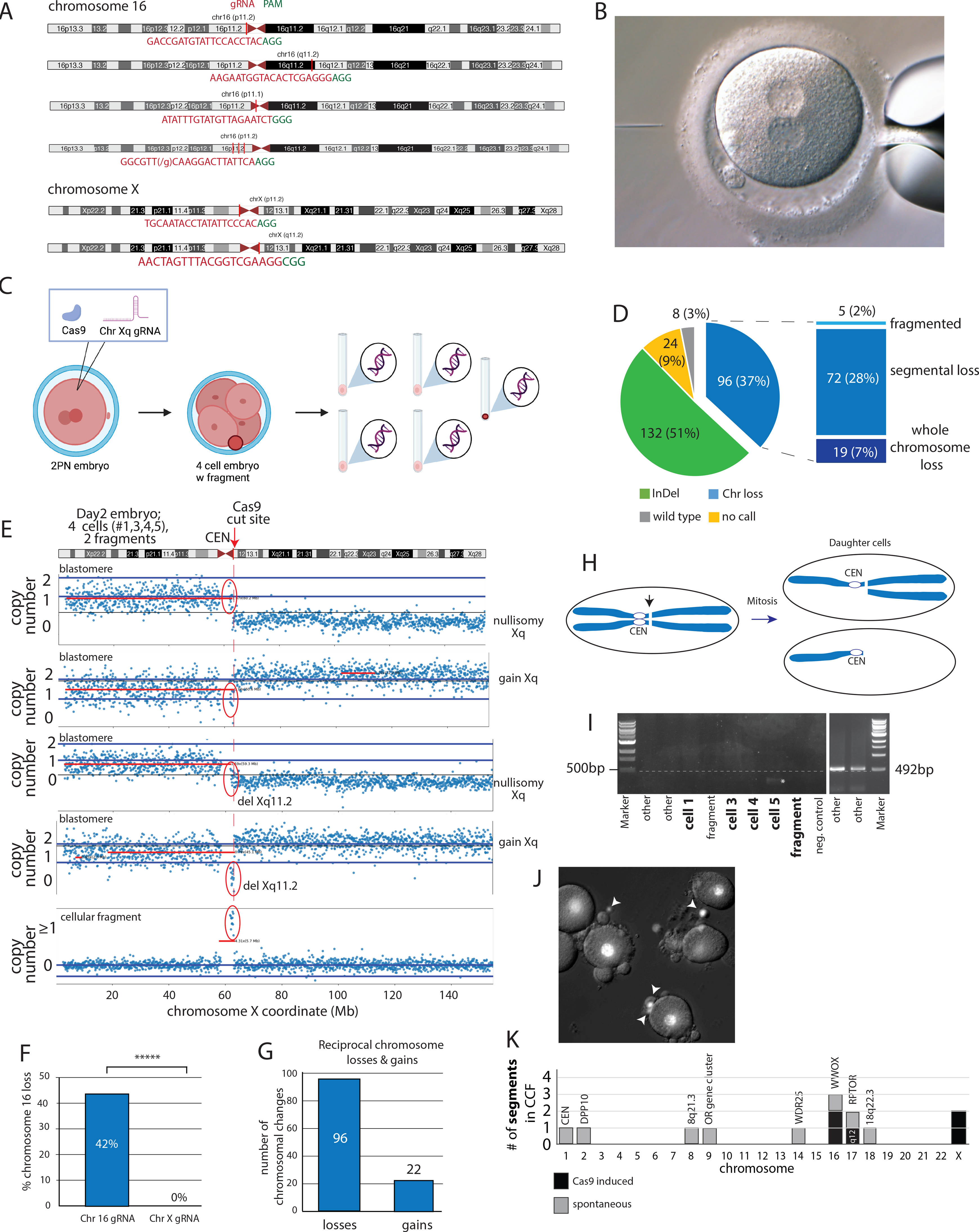
Induction of pericentromeric CRISPR/Cas9 double stand breaks (DSB) frequently result in chromosome loss. a. Guide RNAs to target the pericentromeric region of chromosome 16 on the p arm and the pericentromeric region of chromosome X on the q arm b. Cytoplasmic injection of CRISPR/Cas9 with guide RNA performed on frozen/thawed 2 pronuclear (2PN) human embryos. c. Schematic of the experiment. After CRISPR/Cas9 injection, embryos were cultured for 2 - 6 days followed by single cell collection and analysis. d. CRISPR/Cas9 induced pericentromeric double strand breaks combining results from both chromosomes X and 16 resulted in chromosome loss including segmental, whole and fragmented chromosome loss. Segmental changes are defined as those encompassing a chromosomal arm from the Cas9 cleavage site to the telomere. Results labeled as other had a complete chromosome complement according to SNP array and no discernable genomic sequence on PCR or Sanger sequencing. e. Chromosome X copy number analysis of a day2 embryo (D3_16qXq) consisting of 4 cells, two cell divisions after Cas9 injection. Note the copy number changes of the q arm, with reciprocal losses and gains. f. Chromosomal specificity of Cas9-induced loss. Loss of chromosome 16 was more likely to occur in embryos injected with gRNA targeting chromosome 16 compared to embryos injected with gRNA targeting chromosome X (****p ≤0.00001). g. Quantification of chromosomal losses and gains, combining results from both chromosomes X and 16. h. Schematic for reciprocal loss and gain of acentric chromosome segments in sister blastomeres. A lack of spindle tension results in nondisjunction. i. PCR using primers flanking the Cas9 cut site of the blastomeres of embryo D3_16qXq. *the sequence of this faint band could not be mapped to the human genome and is considered an artifact. j. Day3 embryo stained with Hoechst to identify chromosome-containing cellular fragments. k. Chromosomal segments excluded from the embryo within cytoplasmic cellular fragments. Break sites of spontaneous breaks are indicated with coordinates provided in Table S4.

Embryos were then cultured to the cleavage stage. One hundred and sixty-nine individual cells or cellular fragments from 29 embryos were harvested and individually studied **(Fig 1c, Table S2, Table S3)**. Embryos were analyzed at the cleavage stage on day 2 or on day 3, and a single embryo was a blastocyst on day 6 of development. Cellular fragments, defined as no nucleus in bright field microscopy and smaller than nucleated blastomeres in the same embryo, were analyzed as well, as a previous study showed they could contain excluded chromosomes ^2^. DNA from each sample was amplified and analyzed using a high-throughput SNP array which probes over 800,000 genomic loci to determine copy number and allele heterozygosity along chromosomal arms ^19^. End joining and large deletions were evaluated through PCR and Sanger sequencing. Of the 169 samples, 6 samples were chromosome-containing cellular fragments. Ten samples from two embryos revealed loss of heterozygosity along all chromosomes, which is indicative of haploidy or uniparental diploidy embryos and were excluded from analysis (**Table S3**). Eighteen individual samples showed no detectable genomic DNA, which may be due to being a cellular fragment or due to technical causes (**Table S3**) and 8 cells showed complex chromosomal changes that were not informative with regard to Cas9 mediated aneuploidy. In total, 108 cells and 5 chromosome containing cytoplasmic fragments from 27 embryos allowed evaluation of the consequences of Cas9 activity on the targeted chromosomes **(Table S2)**.

Since blastomeres are diploid, each blastomere contains two homologous chromosome 16s and either one X chromosome in male embryos (XY) or two homologous X chromosomes in female embryos (XX). The gRNAs for chr16p and chrXq were not designed to be allele specific, thus targeting both homologous chromosomes. Among a combined 260 paternal and maternal chromosomes targeted, 37% (96/260) demonstrated a whole, segmental or fragmented chromosome loss (19/260 [7%] whole chromosome loss; 72/260 [28%] segmental chromosome loss; 5/260 [2%] fragmented chromosome loss **(Fig 1d)**. Chromosomal changes were evidenced by both reduced copy number as well as loss of heterozygosity on SNP-array based chromosome screening (**Fig. S1a)**. Abnormal SNP patterns on either side of the targeted locus indicated segmental chromosome abnormalities (**Fig. S1a-c, e, f**) and a whole chromosome loss was detected if reduced copy number and loss of heterozygosity spanned both chromosome arms (**Fig. S1d**). Fragmented chromosomes were evidenced by sporadic chromosome segments amplified and detected by copy number (**Fig. S1g**).

Chromosome losses also occur spontaneously. For example, even after excluding the targeted chromosome 16 and X, over half the cells analyzed from cleavage stage embryos demonstrated aneuploidy for at least one chromosome (**Table S2**). The break points in the observed chromosome 16 and chromosome X segmental losses were noted to occur at the Cas9 cut site; thus, segmental losses can be attributed to CRISPR/Cas9 activity and are not spontaneous **(Fig. 1e)**. Spontaneous versus Cas-9 induced whole chromosome losses cannot be molecularly distinguished. Chromosome 16 losses were compared among embryos injected with gRNA targeting chromosome 16 versus embryos injected with gRNA targeting chromosome X. Loss of chromosome 16 was significantly more likely to occur when embryos were injected with gRNA targeting chromosome 16 compared to embryos injected with gRNA targeting chromosome X (p ≤ 0.00001, **Fig 1f**).

Among the 260 chromosomes targeted, 96 chromosome losses and 22 chromosome gains were identified. **(Fig 1g)**. This represents ~4.5-fold more chromosome losses compared to chromosome gains for the targeted chromosomes. Embryos with a chromosomal loss for the targeted chromosome in one cell often demonstrated a reciprocal chromosome gain in another cell. For example, four cells from one male embryo injected with gRNA targeting the pericentromeric chromosome X q arm were collected on day 2 of development. Two of the four blastomeres showed nullisomy for chromosome Xq; the other two blastomeres had a chromosomal gain of acentric X q (**Fig.1e**). Both segmental gains in this embryo were acentric chromosomal arms. This is consistent with studies in yeast, which show that a single DSB results in co-segregation of acentric chromosome arms ^20^ (**Fig. 1h**). Two cells from this same embryo also showed nullisomy for the chromosomal material Xq11.2 distal to the Cas9 cut site, without corresponding gain in a sister blastomeres **(Fig 1e)**. This represents loss of chromosomal material between the cut site and the centromere. The lost pericentromeric DNA of chromosome Xp11.2 was found in a cellular fragment containing no other genomic DNA (**Fig. 1e**). Such events arise from fusion of sister chromatids of the centromere-containing arms, resulting in dicentric chromosomes, bridge formation and breakage during mitosis ^21^. None of the cells showed specific amplification products with PCR primers flanking the Cas9 cut site (**Fig. 1i**). Though no chromosomal arms had yet been lost in this embryo, all Xq and Xp arms were either disjointed or joined abnormally.

We stained 7 embryos treated with Cas9 with Hoechst and found that 6/7 (86%) showed excluded cytoplasmic fragments containing DNA (**Fig. 1j**). Whenever possible, we sequenced cytoplasmic fragments separately. For 5 of 27 embryos, we amplified and sequenced chromosomal segments contained in cytoplasmic cellular fragments. These showed genomic coordinates of break points consistent with Cas9 cleavage on chromosomes 16 or X (**Fig. 1k, Table S4**). In addition, chromosomal segments from one site on chromosome 17 was also seen, which is consistent with a secondary target site of the guide RNA (**Fig. S2**). No other genomic DNA was detected in these cytoplasmic fragments. Thus, exclusion of chromosomal material from blastomeres in cytoplasmic cellular fragments represents a primary mechanism of chromosome loss from the embryo, contributing to the excess of Cas9-mediated losses over gains in blastomeres. Chromosomal segments resulting from spontaneous breakages were also observed in cellular fragments, all of which were unique in the cohort of embryos. Breakpoints at spontaneous break sites are located at WWOX, an olfactory receptor (OR) gene cluster, WDR25, DPP10, RPTOR, two intergenic sites and the centromere of chromosome 1 (**Fig. 1k**). None of these sites were in proximity to a projected off-target site.

Overall, among the 27 embryos injected with CRISPR/Cas9 and gRNA targeting the pericentromeric region of a chromosome 16 or chromosome X, 24 (24/27, 89%) demonstrated an unrepaired or misrepaired chromosomal loss in at least one blastomere. These results add to the growing body of literature demonstrating frequent and specific chromosome loss after CRISPR/Cas9. For acentric fragments, chromosome loss occurs due to the lack of a centromere. For centromere-containing chromosome segments, a role of bridge formation may play a role, analogous to what has been observed in cultured human cells ^21^.

### Chromosome Removal in Trisomy Embryo by CRISPR/Cas9 DSB

Aneuploidy is exceedingly common in human embryos, occurring in an estimated 20 – 40% of all conceptions^22^. It is the most common cause of pregnancy loss and congenital defects and has become the leading obstacle to the treatment of infertility. Most trisomies arise from errors during maternal meiosis^11^. In this experiment, polar bodies of 10 embryos were collected at the time of CRISPR/Cas9 injection and successfully analyzed by SNP array to identify maternal meiotic errors (**Fig. 2a**). The polar body from one embryo (B2) showed a nullisomy for chromosome 16 **(Fig. 2b)**. This maternal meiotic error is expected to result in a trisomy 16 in all embryonic cells. Interestingly, the same embryo showed reciprocal segregation errors of chromosome 13: the first polar body showed a loss in chromosome 13 genomic DNA and the second polar body showed a gain in chromosome 13 genomic DNA (**Fig. 2b**). This resulted in a normal chromosome 13 complement in the embryo (**Fig. 2c**), a spontaneous correction similar to what has previously been observed ^23^. Thus, information of both polar bodies is required to infer maternal aneuploidies in the fertilized zygote. This embryo (B2) was injected with CRISPR/Cas9 targeting the pericentromeric p arm of chromosome 16 at the pronuclear stage. The embryo was then cultured to day 2 at the 4-cell stage. All 4 cells were analyzed with SNP array, which demonstrated copy number changes for chromosome 16, and PCR analysis revealed indel formation at the targeted sites (**Table S2**). One cell demonstrated, as expected, 3 copies of chromosome 16. Two cells showed a numerical imbalance of 16p and 16q arm (**Fig. 2d**). The remaining fourth cell showed 2 balanced copies of chromosome 16 (**Fig 2d**, bottom). This embryo provides the first evidence that CRIPSR/Cas9 can remove a trisomy in the preimplantation human embryo. However, chromosomal changes were also observed on chromosome 17, as a perfect secondary gRNA target sequence is also found at chr17q12 (**Table S2**, **Fig. 2c**). No further development was tested and thus it is not known whether this embryo would be viable.

**Figure 2.**
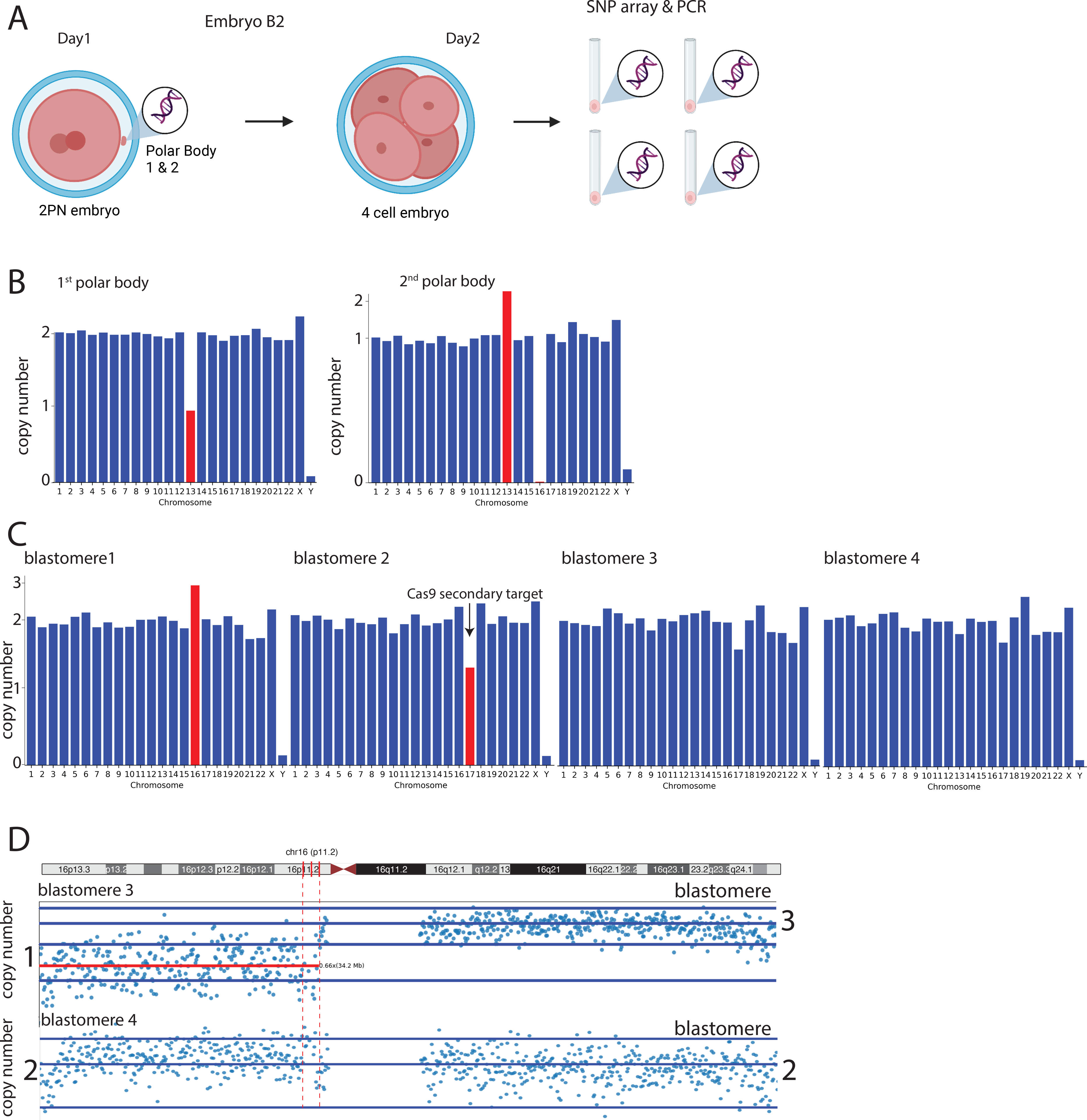
Mosaic correction of a trisomy 16 embryo. a. Schematic of the experiment. Fertilized zygotes have polar body 1 and polar body 2 removed for genotyping. b. Copy number analysis across the genome as a bar diagram. Note the loss of one chromosome 13 in polar body 1, and and the gain in polar body2, resulting in a normal chromosome 13 complement in the embryo. In contrast, chromosome 16 is lost in polar body 2, resulting in a gain in the embryo. c. Copy number analysis across the genome as a bar diagram of 4 embryo cells on day2 of development. d. Copy number analysis across chromosomal arms in two cells. One cell is normal, while the other shows imbalanced copies of 16q and 16p arm.

Previous studies have shown that the use of multiple guide RNAs could result in frequent Y chromosome loss in mouse embryonic stem cells and zygotes^24^. To determine whether multiple gRNA targeting one chromosome resulted in more chromosome loss, 5 embryos were injected with 3 guide RNAs targeting chromosome 16 at 3 different sites: the pericentromeric p arm, centromere and pericentromeric q arm of Chromosome 16. For comparison, 10 embryos were injected with gRNA targeting a single site only at the pericentromeric p or q region of Chromosome 16. When specifically looking at the rates of chromosome loss, they were similar between embryos injected with one vs. three gRNAs (1 gRNA: 43% [31/72] vs 3 gRNA: 43% [25/58]).When considering all chromosomal changes, including losses and gains on chromosome 16, 1 single gRNA site resulted in 66% (24/36) of blastomeres with chromosomal 16 copy number changes. Three separate gRNAs on chromosome 16, of which one gRNA was affected by a common SNP, resulted in 86% (25/29) of blastomeres with chromosomal 16 copy number changes. The difference in chromosomal 16 copy number changes per blastomere was not statistically significant whether one or three cut sites were used (66% [24/36] vs 86% [25/29], p = 0.0872). These results show that chromosome loss can be the most common outcome of a Cas9 induced DSB in human embryos, and can be very high when several gRNAs are combined. Chromosome removal may be increased further than with 3 gRNAs, as the gRNA targeting the centromeric site had a common SNP close to the PAM site that affected gRNA function and also prevented the formation of indels (Table S2).

### CRISPR-Induced DNA DSB may be used to map the functional chromosome centromere

Injection with CRISPR/Cas9 and gRNA targeting the pericentromeric region of the chromosome 16 arms resulted in a large number of segmental chromosome losses (65 segmental losses among 180 targeted chromosomes). The majority of these segmental losses affected the q arm (53 del 16q vs. 12 del 16p). Surprisingly, when injected with a gRNA targeting the chromosome 16 p arm (gRNA Chr16:35126545, at 16p11.1) the majority of chromosomal losses involved the chromosome 16 q arm (1 whole chromosome loss, 1 segmental p loss, and 11 segmental q loss among 30 chromosome targets) **(Fig 3a-d)**. The gRNA targeting the p arm at 16p11.1 was designed using genomic sequences near the chromosome 16 centromere as annotated in the UCSC genome browser (chr16:36-38.5Mb, hg38). According to the annotation of the human genome (hg19 and hg38), the q arm would be linked to a centromere at chr16:35.2-38.5Mb, and therefore be able to segregate to daughter cells. In contrast, the p arms would have no centromere according to hg19 or hg38, and thus no means to reliably segregate sister chromatids to daughter cells (**Fig. 3e**). Furthermore, gain of 16q was observed with reciprocal loss in sister blastomeres (**Fig. 3a**), suggesting nondisjunction and absence of a functional centromere distal of the gRNA cut site. No other site targeted by Cas9 in this or in a previous study acted in this manner. All gains at 8 other sites were found to be acentric chromosome segments, and losses were more common for acentric arms than arms containing a centromere. This pattern of acentric gains, as well as acentric losses exceeding centric losses is consistent with prior studies in yeast cells ^20^. Thus, the annotated chr16 centromere is not functional in human embryos. Approximately 2 MB proximal to the annotated chromosome 16 centromere, a genomic sequence of 0.7 MB also contains alpha satellite DNA repeats (Chr16:34219584 – 34939040, hg38), characteristic of a centromere’s DNA sequence (**Fig. 3b**). If this site acted as the functional centromere, the pattern of chromosome loss would depend on whether cleavage occurs distal or proximal to this region, rather than whether it occurs distal or proximal to the annotated centromere (**Fig. 3f**).

**Fig. 3.**
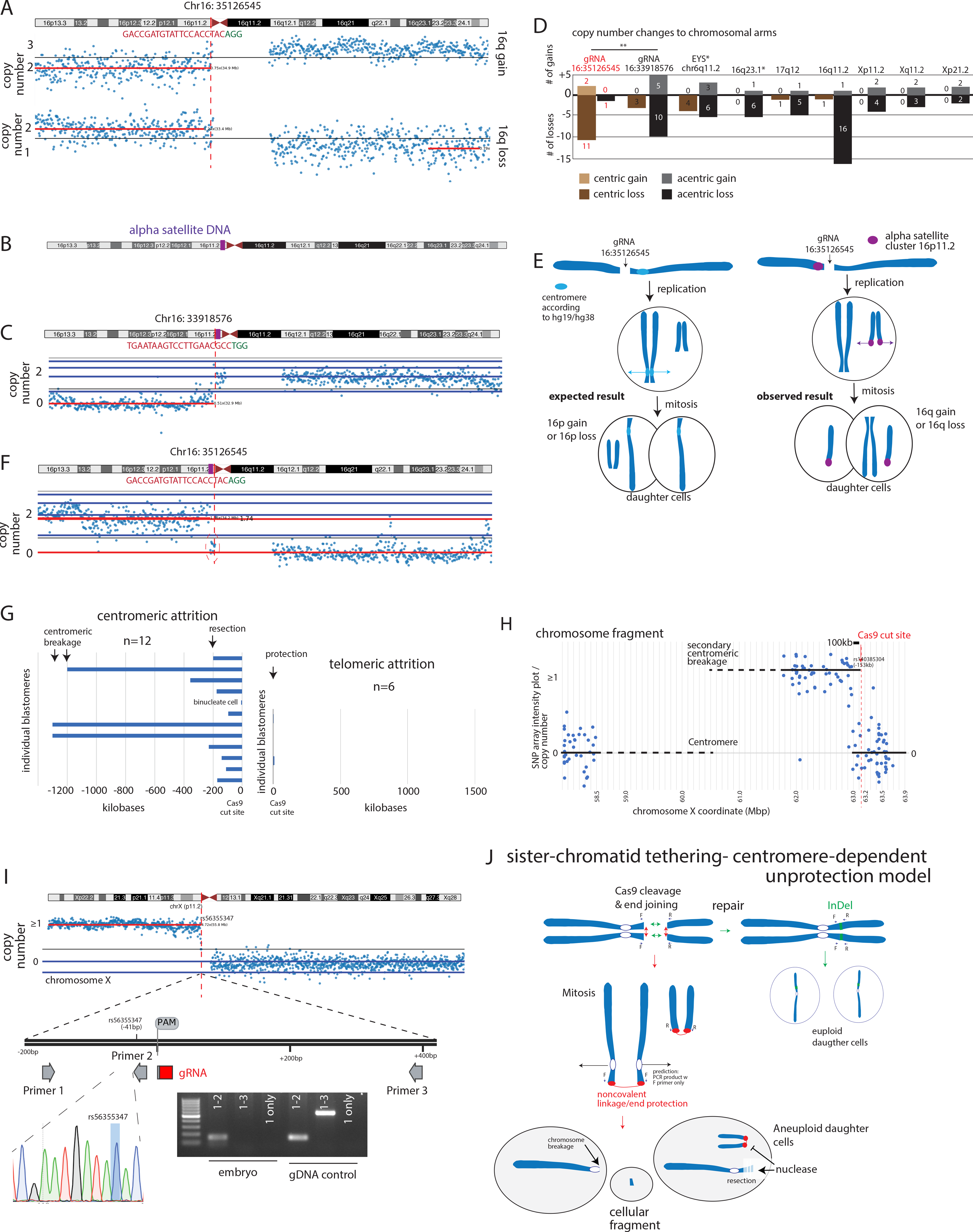
Asymmetric outcomes of Cas9 cleavage relative to the centromere. a. gRNA targeting chromosome 16 at Chr:16:35126545 (hg38). Copy number plot of two blastomeres from the same embryo after targeting of this gRNA. Note the gain of chromosome 16q in one of the two blastomeres and reciprocal loss in the other. b. Approximately 2 MB proximal, a genomic sequence of 700,000 bp contains alpha satellite DNA repeats (purple box on Chr16: 34219584 – 34939040). c. gRNA was designed proximal to this sequence (Chr16:33918576). Copy number plot of blastomere after targeting of this gRNA. Note the loss of the p arm at the cut site. d. Quantification of segmental chromosome gains and losses depending on centromere location ** p< 0.01. Fishers exact test. All sites except gRNA at chr16:35Mb show gains of only acentric arms, and more losses on acentric arms as opposed to centromere containing segments. The pattern meets expectations with a gRNA designed to cut telomeric of alpha satellite DNA located at chr16:33.9Mb. e. Model for chromosome segregation errors after Cas9 cleavage. Chromosome 16 contains two potential regions that might act as a centromere: the annotated region (blue circle), or alpha satellite DNA on 16p11.2 (purple). Cleavage proximal to the annotated centromere would result in an acentric p arm, resulting predominantly in 16p losses and gains. If the purple satellite DNA at 16p11.2 forms the active centromere and the annotated centromere is inactive, gains and losses of an acentric 16q would be seen predominantly. f. gRNA targeting chromosome 16 at Chr:16:35126545. Copy number plot of blastomere after targeting of this gRNA. Note loss of chromosomal material proximal to the cut site (circled), proximal to the active centromere of chromosome 16. g. Asymmetric loss of chromosomal material relative to the Cas9 cut site. Chromosomal breaks due a single cut site on chromosomes 16, 17 and X were analyzed. Genomic coordinates of chromosome copy number transitions are annotated in Table S4. h. Example of attrition of gDNA centromeric to the Cas9 cut site. A secondary centromeric breakage is also observed. i. Conservation of the break site telomeric of the Cas9 cut site. PCR primers telomeric to the break site amplify gDNA, and signal of a SNP adjacent to the cut site is detected. j. Model for the loss of pericentromeric material centromeric to the cut site. Sister chromatids are tethered at broken DNA ends through protein-mediated linkage, which act antagonistically to spindle forces centromeric but not telomeric to the Cas9 cut site. This model is consistent with both the asymmetry of chromosomal changes as well as with the asymmetry of attrition at the break site.

To test if cleavage telomeric to Chr16:34219584 – 34939040 alters the pattern of chromosome loss, a new gRNA was designed proximal to this sequence (Chr16:33918576, Table S1) (**Fig. 3c)**. Significantly more segmental p chromosome loss was seen with the new proximal chromosome 16 p arm gRNA (20% [10/50] vs 3% [1/30], p = 0.046) **(Fig. 3d)**. Gains of acentric chromosome 16p were also obtained (**Fig. 3d**). This suggests that the Chr16:34219584 – 34939040 region acts as the functional centromere in human embryos. Previous literature has shown in rare cases, neocentromeres can form at new sites on a chromosome as a result of a repositioning of the centromere and inherited within a family ^25^. Both the original and new chromosome 16p gRNA were injected into embryos from unrelated families with similar results and thus this centromere is commonly active, rather than a neocentromere unique to a specific family. Given the small numbers of samples, more work is still needed to determine whether the annotated centromere can also be active, including in somatic cells. Our results show that DNA DSB induced by CRISPR/Cas9 may be used to determine the location of the functional centromere in human embryos based on the segregation patterns of acentric and centric chromosome arms (**Fig. 3e,f**).

### Asymmetric attrition of genomic DNA at the Cas9 break site towards the centromere but not the telomere

We noted that cleavage on chromosome 16p11.2 distal to the functional centromere Chr16:34219584 – 34939040 (hg38), but proximal to the centromere annotated in the genome browser (chr16:36-38.5Mb) resulted in copy number changes distal to the cut site (**Fig. 3g**, **Fig. S3**). Similar loss was also observed on chromosome X (**Fig. 1e**). Such copy number changes may arise from covalent or noncovalent linkage of sister chromatids and breakage of dicentric chromosomes at the centromere.

We mapped loss of genomic DNA in nullisomic copy number transitions with an accuracy of 41bp-22kb using SNP intensity array data, depending on the location of the SNP closest to the Cas9 cut site (**Table S4**). Loss of gDNA was readily apparent in 11/12 centromeric to the Cas9 cut site. Loss of gDNA centromeric to the break site consisted of two forms: breakage at the centromere, resulting in the loss of more than 1 Mb, and attrition of 0.1-0.3Mb (**Fig. 3h**). For instance, one embryo, containing a chromosome X fragment, the closest detectable SNP was rs140385304, 153kb centromeric to the Cas9 cut site (**Fig. 3i**). In addition, this fragment also had a break at the centromere. The only cell with a centromeric fragment but without detectable attrition towards the centromere was binucleate (I1-4), caused by cytokinesis failure.

In contrast, no detectable attrition was observed in 5/6 sites telomeric to the cut site. In one sample, attrition was detectable though smaller than 4.9kb (**Fig. 3h**). We also used PCR primers adjacent to the cut site to narrow attrition of DNA telomeric to the cut site. In a cell containing an acentric chromosome X arm, a primer pair 20bp distal to the cut site resulted in specific amplification, verified by Sanger sequencing (**Fig. 3i**), with no PCR product obtained using primers flanking the cut site. Absence of detectable product using primer 1 alone argues against covalent head-to-head joining of sister chromatids. Furthermore, the closest SNP on the array rs56355347 at a mere 41bp telomeric of the Cas9 cut site showed detectable signal (**Fig. 3i**). In this male embryo (H1), all 8 cells showed reciprocal whole and segmental chromosome changes, demonstrating that the cut site was preserved through 3 cell divisions, for a total of 3 days (**Table S2**).

This remarkable preservation of the cut site suggests end protection telomeric to the Cas9 cut site. In contrast, spindle forces acting on the centromeric side result in attrition at the cut site, as well as in secondary breakages at the centromere. One possible interpretation of these results are head to head ligation of sister chromatids instead of ligation with the telomeric segment of the same chromatid (**Fig. 3k**). The resulting dicentric chromosomes would break between the two centromeres, resulting in loss of gDNA in one cell, and a reciprocal tandem duplication in a sister cell, between the break site and the centromere. The fusion of sister chromatids would also conserve genomic DNA at the break site itself, which is inconsistent with the observed attrition (**Fig. 3g**). Reciprocal losses and gains pericentromeric to the break were observed in only one embryo (**Fig. S3**), though conservation of gDNA at the break site could not be verified because of cleavage at multiple sites. These duplications may be tandem duplications due to head to head joining, or small chromosome segments as in **Fig. 1e**. None of the cells in other embryos with chromosomal changes showed evidence of pericentromeric duplications. However, because of the proximity of the Cas9 cleavage site to the centromere, they may be too short to detect.

### Mid-arm DNA DSBs result in chromosomal changes without tandem duplications

Tandem duplications arising from inter-sister chromatid fusions and breakage of dicentric chromosomes during cells division should be readily apparent when the Cas9 cut site is on the chromosomal arm, such as on chromosome 17, located 11Mb from the centromere. We also designed a gRNA to target the dystrophin gene responsible for Duchenne muscular dystrophy located in the short arm of the X chromosome, at Xp21.2, 27Mb from the centromere. Breakage between the cut site and the centromere would result in reciprocal and variably sized tandem duplications and deletions, detectable by copy number analysis. Five embryos were injected with the gRNA targeting the mid p arm of chromosome X and resulted in 12 cells and a total of 17 chromosome X targets. From SNP array analysis, two Xp segmental losses were identified (chromosome loss: 2/17, 12%) **(Fig. S4a)**. Segmental chromosomal changes were also seen with Cas9 cleavage on chromosome 17q12 in 7/24 cells (48 targets, 29%) (**Fig. 3d**). Furthermore, two prior studies with cleavage midarm showed chromosomal changes, on chr6p ^3^ at a frequency of 16% (4/25), and on chr16q23.1 ^2^ at a frequency of 21% (7/33). The frequencies of segmental changes due to cutting at mid-arm was not as common as pericentromeric on chromosome 16 or chromosome 6, but the differences in frequency between chromosomal positions (arm versus pericentromeric) were not statistically significant because of high variation between sites **(Fig. S4b,c)**. No tandem duplications were found in any sample (0/99 chromosomal targets).

Thus, noncovalent tethering rather than end joining of sister chromatids account for the majority of chromosomal changes. Intersister repair cannot be ruled out for all samples, but it is not the primary mechanism of chromosomal aneuploidies induced by Cas9. Linkage counteracts spindle-forces, resulting in the loss and/or breakage of centromere-containing chromosome segments, as well as in unprotection and attrition of chromosome ends, while acentric segments remain linked, resulting in both reciprocal losses and gains as well as protected chromosome ends (**Fig. 3k**).

### The vast majority of end joining events are small deletions and insertions, larger deletions are predominantly mediated by MMEJ

Fifty one percent (132/260) of chromosomes targeted at pericentromeric locations of chromosome X or 16 revealed an indel at the targeted location and 3% (8/260) were found to be wild type. Thus, the pericentromeric gRNAs demonstrated a very high on-target efficiency for the targeted region. Twenty-four samples (9%, 24/260) had a complete chromosome complement according to SNP analysis but unknown genomic sequence at the targeted DNA DSB due to lack of a specific PCR product. These may represent unrepaired or misrepaired chromosomes, or technical failure.

To assess repair outcomes through end joining at each targeted region, 500 base-pair (bp) PCR primers that span the gRNA target sites were used followed by Sanger sequencing. In total 96 sites targeted by gRNA resulted in no PCR band with confirmed chromosome loss on SNP array analysis (37%, 96/260). Through Sanger sequencing analysis, 132 sites (51%, 132/260) revealed an indel at the targeted location.By far the most frequent indel among all samples in our experiment using human embryos was −1 bp deletion and a +1 bp insertion, which may be the result of processing the stagger created by Cas9 cleavage ^26^ **(Fig 4a)**. Gel electrophoresis showed smaller PCR DNA fragments **(Fig. 4b)** that were confirmed by Sanger sequencing to be the result of CRISPR/Cas9 induced deletions **(Fig. 4a)**. Indels ranged from −358bp to +2bp **(Fig. 4a)**. To assess for larger deletions, 1.3- kb PCR with primers equidistant from the gRNA sequence on chromosome 16 p arm was also used. Only 3% of chromosome targets at this location showed a deletion more than 100 bp (n = 1 among 30 chromosome targets). Among 157 indels at all sites, only 3 were larger than 100bp (**Fig. 4a**). In contrast, a prior study in mice found deletions greater than 100 bp in over 44% of samples (57/127)^27^. This significant difference (p=0.0007, Fishers exact test) in the frequency of large deletions in a comparable assay between mouse and human embryos appears to be consistent across sites: other studies in human embryos have also found few or no large deletions ^9^. Deletions of 6 or more base pairs in human embryos occurred primarily through microhomology-mediated end joining (MMEJ) (**Fig. 4c**). Microhomology-mediated end joining uses short sequences of homology and results in the deletion of the base pairs in between, while nonhomologous end (NHEJ) joining does not.

**Fig 4.**
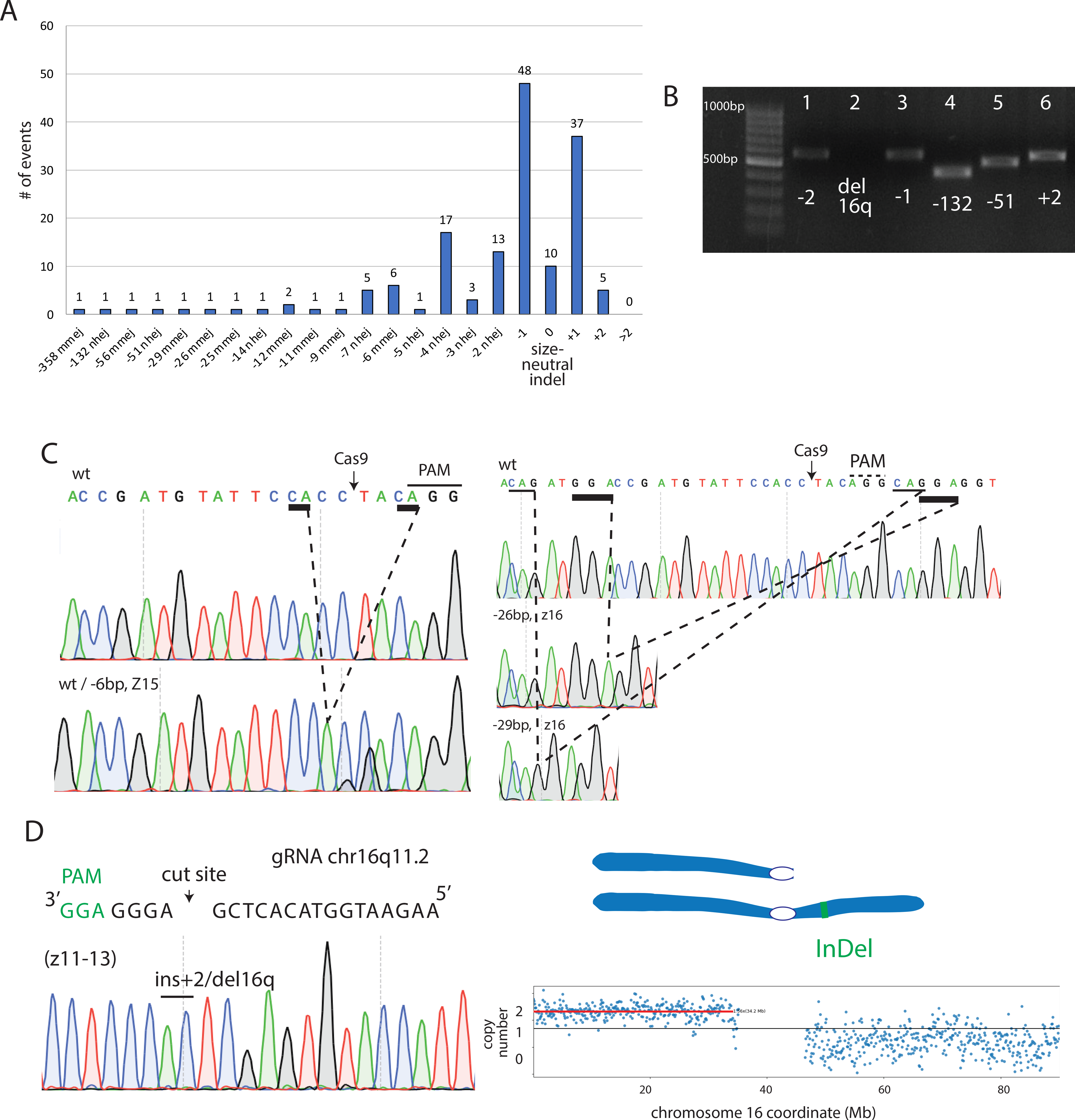
Large indels are uncommon in human embryos. a. Quantification of indel frequencies at the pericentromeric targeted locations on chromosome 16 and chromosome X. Of note, embryos injected with gRNA targeting chromosome 16 at multiple locations (Chr 16 p arm, centromere and q arm) may result in multiple indels at the different targeted locations on chromosome 16. MMEJ = microhomology-mediated end joining events, NHEJ = nonhomologous end joining. b. Gel electrophoresis demonstrating representative examples of different PCR outcomes at the chromosome 16q location after CRISPR/Cas9 and chromosome 16 gRNA injection. Different sizes of PCR DNA products suggest indel formation. If the PCR product failed to amplify as seen in sample 2, this suggests chromosomal changes incompatible with PCR amplification using primers flanking the cut site. Cell IDs from 3 different embryos: 18_Z12, 20_Z13, 16_Z12, 15_Z12, 14_Z11, 13_Z11 (Table S2). c. Examples of microhomology-mediated end joining (MMEJ) at gRNA site chr16:35126552 in zygote 15 (z15) and zygote 16 (z16). Regions of microhomology are underlined and the combination pattern indicated with dotted lines. d. Analysis of a blastomere by Sanger sequencing, revealing a single indel with an insertion of 2 nucleotides (AC). Chromosomal constitution of the same cell by SNP array probe intensity plot shows loss of the q arm. The indel therefore is hemizygous.

Indels were found to be either heterozygous resulting in chromatograms with overlayed peaks, or only a single indel was found. Events with a single indel commonly also showed chromosomal change (**Fig. 4d**). Among all samples, 39 pairs of homologous chromosomes targeted by gRNA were complete according to SNP analysis. Among these 39 pairs, 19 demonstrated different indels or a combination of indel and wild-type at the targeted location. Twenty pairs of homologous chromosomes (51%, 20/39) detected by SNP array had a single indel at the targeted location. These may be two identical indels due to preferred NHEJ indel outcomes, or repair by gene conversion where the homologous chromosome is used as a template. Alternatively, PCR does not detect the presence of an unrepaired break in one of two chromosomes, or abnormally joined chromatids. Heterozygosity within the PCR product would conclusively demonstrate the detection of two chromosomes within a single PCR product. However, no heterozygosity was observed in any of the single indels. Thus, whether these indel events represent one or two chromosome copies remains unknown.

## Discussion

In conclusion, CRISPR-induced pericentromeric DNA DSB resulted in chromosomal loss in over one third of chromosomes targeted. The large number of chromosome losses and gains observed in this study adds to the safety concerns for the future of point mutation editing using CRISPR/Cas9. Both whole chromosome and segmental chromosome aneuploidy could have detrimental consequences on the developing embryo. While the impact of segmental aneuploidies in the preimplantation embryo remains poorly understood, whole or segmental aneuploidies seen at later developmental stages are typically incompatible with life. The only mitotically acquired aneuploidy compatible with development to term is the mosaic loss of the X-chromosome associated with Turner syndrome ^28^. While the human embryo has at least some ability to exclude cells with mitotically acquired aneuploidies ^29^, attempts to edit genes on the X chromosome such as DMD using Cas9 would carry a high risk of congenital abnormalities.

This study reveals new insight into the mechanisms of chromosome loss after Cas9 cleavage in human embryos. While chromosomal losses were observed in both the acentric and to a lesser extent also in centric chromosome segments created by Cas9 cleavage, only acentric fragments produced chromosomal gains. This notion holds true for all sites tested, including for chromosome 16 after correcting centromere location to an area currently annotated as chr16p11.2. The nonrandom allocation of sister chromatids to daughter cells suggests linkage of sister chromatids persisting through mitosis.

Asymmetry was also seen in the attrition of genetic material relative to the Cas9 cut site. Centromeric of the Cas9 cut site, we observed attrition of several hundred kb, as well as secondary breakage at the centromere. Only one of twelve instances showed no attrition centromeric to the Cas9 cut site, but this cell had failed cytokinesis, containing two nuclei. Pericentromeric losses of genomic DNA adjacent to the cut site suggests that sister chromatids are linked, followed by rupture of that link during mitosis. DNA bridge formation has been observed in Cas9-mediated chromosomal aneuploidies in mouse embryos ^30^ as well as in cultured cells ^21^. MRE11 forms dimers binding to DSB ends, thereby tethering them ^31^. In yeast cells, broken chromosome ends are tethered through protein-mediated linkage involving MRE11, and unrepaired breaks can be passed through mitosis without covalent linkage ^20^. One of MRE11’s numerous functions is to prevent the conversion of a DSB into a chromosome break by tethering DSB ends in cis ^32^. However, MRE11 complexes can also directly contribute to aneuploidies by tethering sister chromatids. Tethering a chromosome break to the sister chromatid occurs during the repair of one-ended DSBs that arise spontaneously during DNA replication at damaged replication forks, which allows use of the sister chromatid as a template for repair ^33^. However, in the context of a two-ended DSBs as generated by Cas9, the sister chromatid is not a suitable repair template. If a replisome encounters an unrepaired DSB, the broken sister chromatids are in close proximity and may hence be tethered by MRE11 (Model in **Fig. 3j**).

On the centromeric side, tethering of broken sister chromatids counter spindle forces, resulting in the trapping of DNA in the midbody, which can result in secondary chromosome breakage through abscission, as well as in the deprotection and attrition of chromosome ends. Alternatively, it may result in cytokinesis failure and binucleation. In contrast, on the telomeric side, acentric fragments are not exposed to spindle forces, and MRE11 complexes may remain bound to chromosome ends, protecting them from extensive degradation. Covalent head-to-head joining of sister chromatids would equally conserve the break site telomeric and centromeric of the Cas9 cleavage site, which is inconsistent with asymmetric attrition and concurrent secondary breakage at the centromere. Mre11 nuclease activity is required for MMEJ ^34^, as well as for HR ^35^. MMEJ was observed in human zygotes in both this and an earlier study ^2^ and thus, we know that Mre11 is binding to and processing Cas9-induced DSB ends at the earliest stages of human embryonic development. Once committed to homologous recombination instead of end joining, the break may remain unrepaired. By tethering unrepaired sister chromatids in mitosis, MRE11 may be a key contributor to aneuploidy, micronucleation and cytokinesis failure in the human embryo.

Prior studies have suggested CRISPR/Cas9 may be used to intentionally eliminate an extra chromosome^36,37^. In a proof-of-concept study, Adikusuma demonstrated efficient CRISPR-mediated selective chromosome deletion by removing the centromere or shredding the chromosome arm with multiple targeted gRNA sites in mouse embryonic stem cells and zygotes ^36^. In order to apply this technology to an aneuploid embryo, the specific trisomy would need to be detected prior to CRISPR/Cas9 injection. Embryos with a potential trisomy due to errors in maternal meiosis could be identified by polar body analyses, which is feasible and compatible with live birth ^23^. While the first polar body can be biopsied at fertilization, the second becomes available within 4-5h after fertilization. Data from McCoy and colleagues demonstrate that approximately 5% of human embryos possessed a single trisomy that could be edited ^38^. In order to avoid uniparental disomy or the loss of two targeted chromosomes after editing, the gRNA of CRISPR/Cas9 would need to be parental-specific and distinguish between the two homologous chromosomes from the same parent. Timing of the CRISPR/Cas9 injection is also crucial. Injection with CRISPR/Cas9 even at the 1-cell stage can result in chromosomal mosaicism as the loss of centric and acentric chromosome segments requires several cell cycles. In this experiment, all embryos injected with CRISPR/Cas9 on day 1 at the 2PN stage demonstrated mosaicism at the cleavage stage. However, a prior study showed that embryos with uniform chromosome loss can result at the blastocyst stage ^2^. Analogous to observations on spontaneous aneuploidies ^29^, at least some of the Cas9-mediated aneuploidies appear to resolve with continued development. Despite these promising advances, an extensive body of basic science research is still needed to determine the timing, efficiency and safety of intentional chromosome elimination to prevent trisomy.

In recent years, preimplantation genetic testing for aneuploidy, which has been used to identify euploid embryos prior to transfer, have also identified segmental aneuploidy at appreciable frequencies ^39–44^. The majority of spontaneous segmental aneuploidies are believed to be mitotic in origin and arise during the first few mitoses following fertilization ^45^. Hence, segmental aneuploidies detected in human cleavage stage embryos may be the result of a spontaneous DNA DSB during the first few mitotic divisions following fertilization. In support of this notion, we found chromosome segments contained in cellular fragments with break points in known fragile sites such as WWOX and DPP10. Both genes are known fragile sites that break spontaneously in tumor cells ^46,47^ and have been implicated in developmental disorders of the nervous system ^48,49^. Chromosome-containing cellular fragments have previously been observed in both human and rhesus macaque embryos ^50,51^, though neither the location of chromosome breakage, nor cause and consequence have previously been experimentally identified. Our studies show that a single DNA DSB is sufficient to induce sequestration of a chromosome in a cellular fragment. Tethering of sister chromatids at spontaneous DSBs may also result in chromosome loss, micronucleation, and cytokinesis failure, which is commonly observed in human embryos. Additional studies will be needed to determine the origin of spontaneous DSBs, and the mechanisms determining DNA repair pathway choice. CRISPR-Cas9 induced DNA DSB provide a model to study DSB repair in the embryo, and provide a tool to study aneuploidy, including its origins, progression, surveillance, and even potential treatment.

### Limitations of study

This study does not include functional interrogation of MRE11 and interacting DNA repair proteins in the human embryo. While there are specific inhibitors for MRE11 nuclease activities, there are not currently any compounds that would interfere with dimerization and DNA end binding. The molecular role of MRE11 in the human embryo is inferred based on the knowledge obtained in yeast and somatic cell lines.

Further limitations of this study include the potential presence of other genetic outcomes from CRISPR/Cas9 DSBs such as complex genomic rearrangements and translocations not seen on SNP-arrays or by PCR genotyping the targeted genomic locus. However, these other outcomes would need to be linked to abnormality of another chromosome, and also do not explain the asymmetry of chromosome losses and gains as well as the asymmetry of attrition relative to the Cas9 cut site.

For the analysis of on-target indels, we were unable to distinguish if identical indels identified on homologous chromosomes were the result of preferred NHEJ outcomes, presence of a cleaved or misrepaired chromosome that is not detected by PCR, or due to interhomolog homologous recombination. Distinguishing these outcomes would require flanking heterozygous SNPs within the same PCR product which conclusively identify recombination between paternal and maternal chromosomes^2^.

And lastly, the CRISPR/Cas9 injection of the gRNA may have also resulted in other off-target effects, including off-target indels, which was not evaluated here.

## METHODS

The Columbia University Institutional Review Board approved all human procedures and experiments.

### Research Samples

#### Human embryos

Cryopreserved human 2 pronuclear (2PN) embryos were anonymously donated from couples who provided informed consent for use in research. Embryos were cryopreserved between the years 2007 and 2012 using One Step (PB1. PG, 1 M sucrose) cryopreservation solution, Sage embryo freeze kit or Quinn’s embryo freeze kit. Embryos were stored in liquid nitrogen until use. For the experimental procedures, embryos were thawed then exposed to experimental conditions as outlined below. All experiments were conducted during incubation at 37°C with 5% CO2 and 20% O2. All human embryos were cultured for no more than 1 – 6 days, in accordance with at the time of conduct of the research internationally accepted standards to limit progression to less than 14 days ^36^.

### Method Details

#### RNP preparation

2nmol of single guide RNAs were obtained for Integrated DNA Technologies (IDT) with the sequence listed in Supplementary Table 1 and dissolved in 20μl to a concentration of 100μM sgRNA. For ribonucleoprotein (RNP) preparation, 3μL of injection buffer, 2μL of 63μM IDT nlsCas9 v3, and 1.5 μL of 100μM sgRNA were combined and incubated at room temperature for 5 minutes. Thereafter, 96.5μL of an injection buffer was added. Injection buffer consists of 5mM Tris-HCl, 0.1mM EDTA, pH 7.8. The RNP solution was then centrifuged at 16000 RCF for 2 minutes prior to loading into the injection needle and cytoplasmic injection.

#### Embryo manipulations

Embryo manipulations were performed in an inverted Olympus IX71 microscope using Narishige micromanipulators on a stage heated to 37°C. Frozen 2PN (day 1) embryos were thawed using One Step or Sage Embryo thaw kit and pronucleus formation was confirmed. Embryo polar bodies were collected whenever possible. RNP was prepared as above. The tip of an injection needle was nicked and small, but visible amounts of the Cas9RNP was injected manually into the cytoplasm of thawed embryos using a Narishige micromanipulator. Embryos were then cultured in Global Total (Cooper Surgical) in an incubator (Thermo Scientific, Heracell 150i) at 6% CO2, 37°C until collection.

#### Genome amplification and genotyping

Single blastomeres were collected on day 2 to day 3, or if indicated, on day 4, on the heated stage of an inverted Olympus IX71 microscope equipped with Narishige micromanipulators and a zona pellucida laser (Hamilton-Thorne). Trophectoderm biopsies were obtained on day 6 of development using 300ms laser pulses to separate trophectoderm from the inner cell mass. All samples were placed in single tubes with 2 or 4 μL of PBS. Amplification was performed using REPLI-g single kit (Qiagen) according to manufacturer’s instructions, using either a half reaction for 2 μL, or a full reaction for 4 μL.

Genotyping was performed using primers for amplification and sequencing as listed in Supplementary Table 5. PCR was performed using AmpliTaq Gold. PCR products were run on a 1.5% agarose gel for visual inspection of product size and submitted to Genewiz for Sanger sequencing. Base changes were analyzed at the region of Cas9 target sites using Snap Gene2 and ICE analysis (Synthego).

#### Genome-wide SNP array

Embryo biopsies were amplified at Columbia University using REPLI-g single cell kit. according to manufacturer’s instructions. Copy number and genotyping analysis was performed using gSUITE software (Genomic Prediction). For copy number analysis, raw intensities from Affymetrix Axiom array are first processed according to the method described (Mayrhofer et al., 2016). After normalizing with a panel of normal males and females, the copy number is then calculated for each probeset. Normalized intensity is displayed. Mapping of endogenous fragile sites was done through visual evaluation of loss of heterozygosity. Break points were mapped to chromosomal bands by visual analysis of SNP array chromosome plots including analysis of both copy number signal and heterozygosity calls. The accuracy of mapping is between 100-500kb. A segmental error was defined as the gain or loss of a chromosome arm or segment.

Fragmented chromosome was called if there were multiple break points within a given chromosome. Chromosomal coordinates were mapped using probe intensity data on samples where chromosomal changes included nullisomy or a difference of at least two copies.

#### Quantification and Statistical Analysis

Statistical analysis was performed using Fisher’s exact test as indicated. A p-value of less than 0.05 was considered significant.

#### Data availability

Data are available at GEO under accession numbers GSE186407. https://www.ncbi.nlm.nih.gov/geo/query/acc.cgi?acc=GSE186407

## Author contribution

J.T. designed experiments, performed molecular analysis of Cas9 cut site, data analysis and interpretation of SNP arrays. D.M., J.X. and N.T. performed SNP array analysis. S.X. performed break point mapping. D.E. provided assistance at all levels of the work. A.R. assisted with embryo thaws. J.T. and D.E. wrote the manuscript with input from all authors.

## Acknowledgments

We thank the Russell Berrie Foundation Program in Cellular Therapies for funding support; Chyuan-Sheng Lin for advice with Cas9 RNP injection. We thank Shuangyi Xu for critical reading of the manuscript. J.T. and A.R. are supported by a clinical fellowship in reproductive endocrinology and infertility. We thank Rodney Rothstein for helpful discussions.

## Conflict of Interest Statement

J.X., D.M. and N.T. are employees and/or shareholders of Genomic Predictions Inc. D.E. and N.T. applied for a patent of chromosome removal.

## Supplemental Figure Legends

**Supplementary Figure 1.**
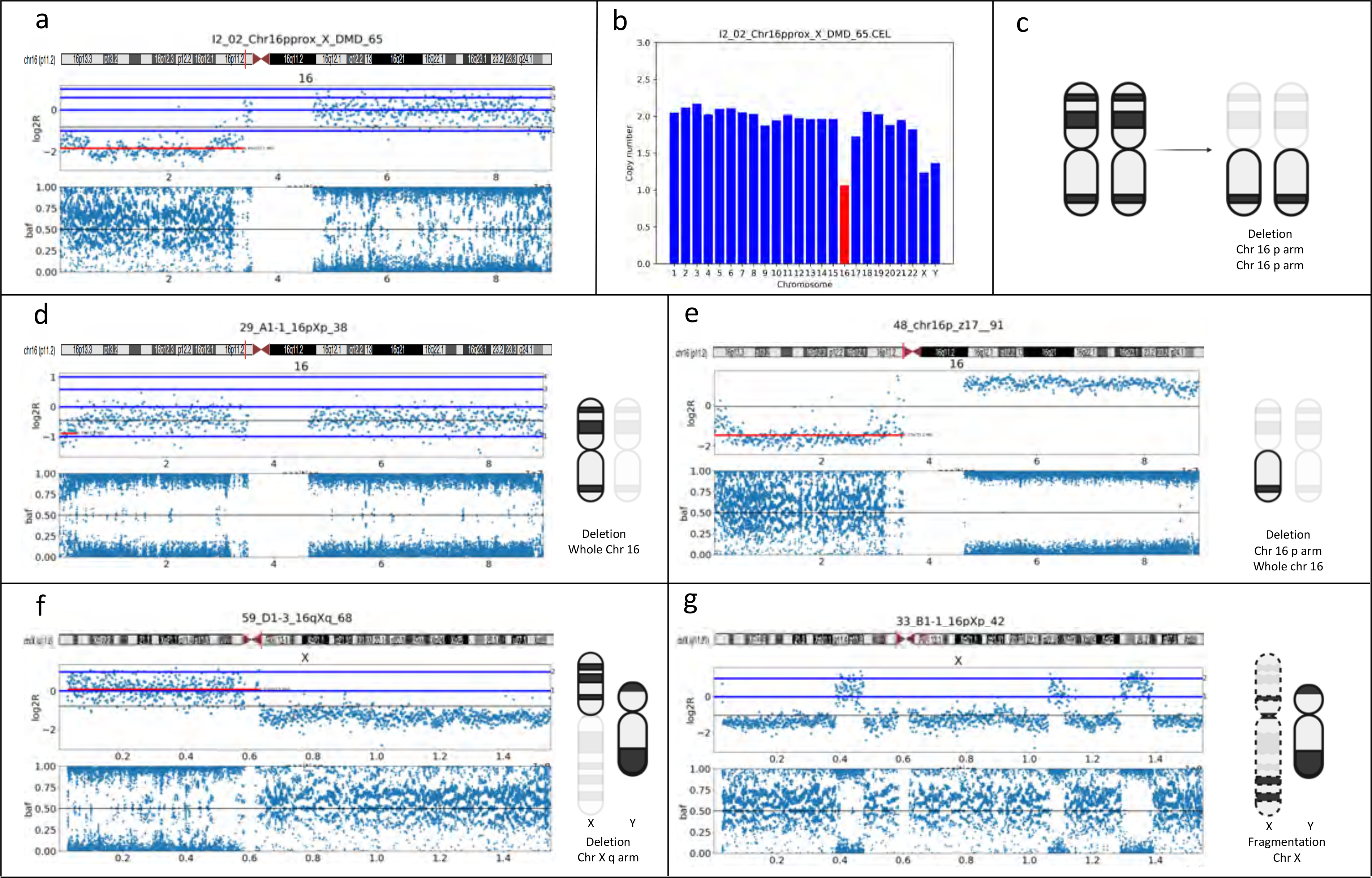
SNP analysis demonstrating CRISPR/Cas9-induced segmental and whole chromosome loss as well as chromosome fragmentation. Representative examples of chromosomal outcomes after targeting chromosome 16 or chromosome X using Cas9. a. After injection with CRISPR/Cas9 targeting pericentromeric region of chromosome 16 p arm, SNP analysis demonstrated complete loss of genomic DNA on chromosome 16 p arms in one cell. Log2R indicates signal intensity of SNP probes. baf indicates B allele frequency. Signals at 1 or 0 indicate only one allele is being detected, while signals in between indicate two are detected, which is heterozygosity. When signals at 0 and 1 are missing, it is a nullisomy and blue dots are noise as here on chromosome 16p. b. The copy number bar graph from the same cell demonstrated loss of genomic DNA on chromosome 16. c. These results indicated segmental loss of both chromosome 16 p arms in this single cell. d. This SNP analysis demonstrated whole chromosome 16 loss due to the loss of genomic DNA in the chromosome 16 p and q arms as well as loss of heterozygosity throughout the chromosome. e. This SNP analysis demonstrated whole chromosome 16 loss and additional segmental loss of chromosome 16 p arm due to the loss of chromosome 16 p arm genomic DNA and loss of heterozygosity on chromosome 16 q arm. f. After injection with CRISPR/Cas9 targeting the pericentromeric region of chromosome X, SNP analysis revealed a segmental loss of chromosome X q arm in this single cell. g. This SNP analysis demonstrated a fragmented chromosome X.

**Supplementary Figure 2.**
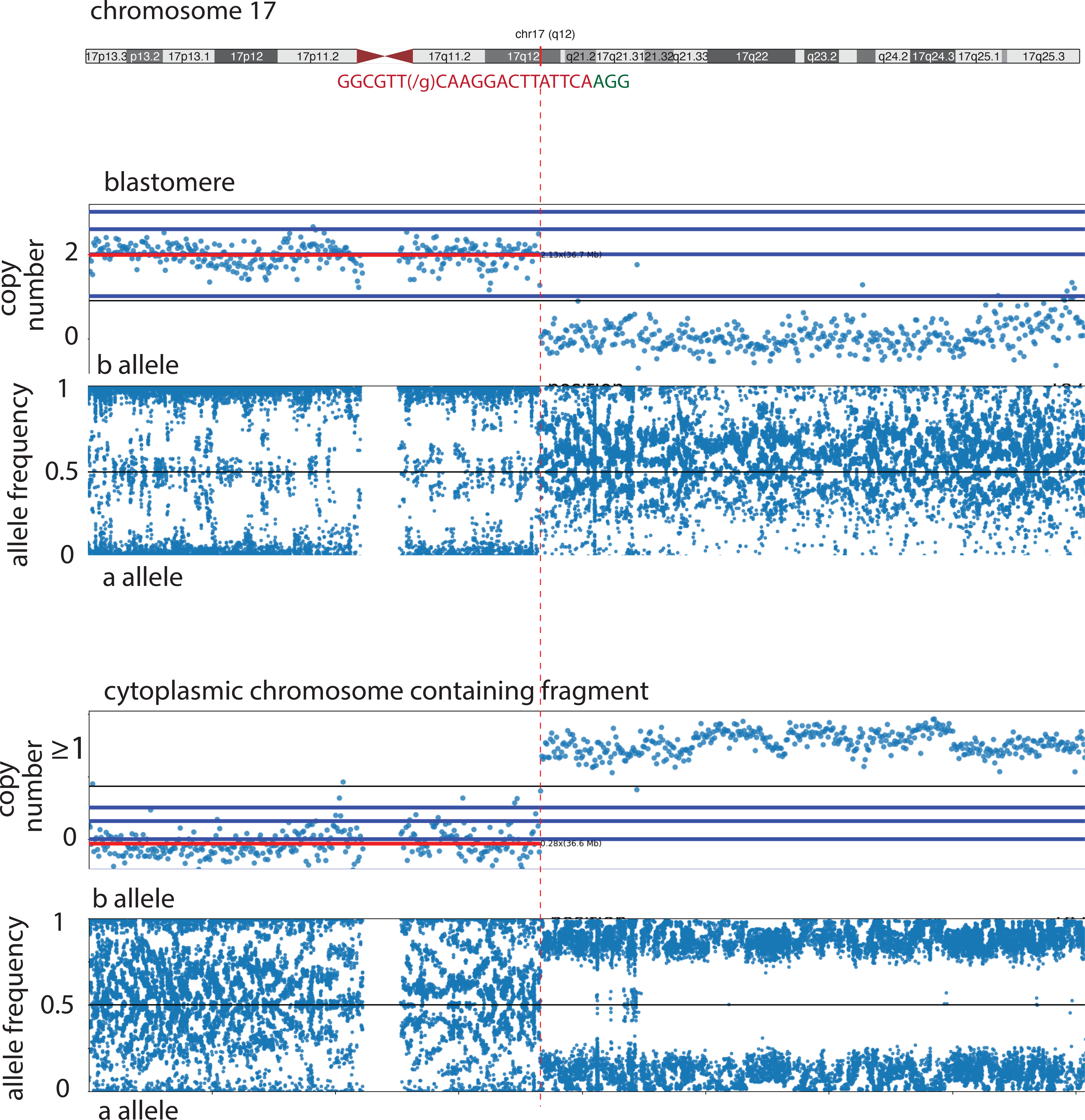
Exclusion of chromosomal segment in a cytoplasmic chromosome containing fragment. Reciprocal chromosome breakage in a blastomere (top) and a chromosome segment excluded in a cytoplasmic fragment (bottom).

**Supplementary Figure 3.**
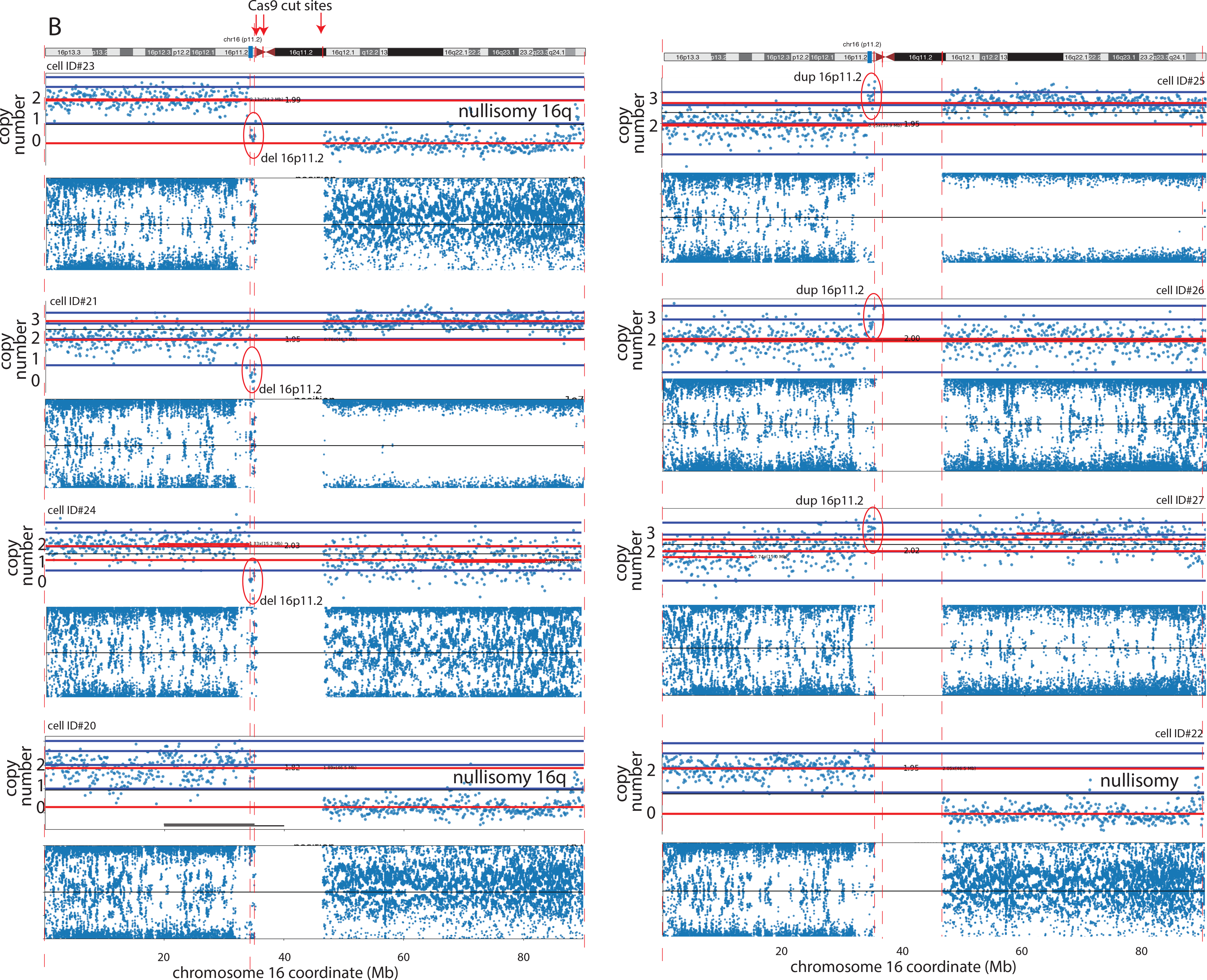
Pericentromeric cleavage results in copy number changes between Cas9 cut site and the centromere. SNP analysis for a 2PN embryo (z13) injected with three gRNAs targeting 3 locations on chromosome 16 (pericentromeric p arm, centromere and pericentromeric q arm). The embryo was injected with CRISPR/Cas9 and gRNA at the 2 pronuclear (2PN) stage and cultured to a day3 cleavage stage embryo; eight cells were individually collected and analyzed. On SNP analysis, red represents chromosome 16 p arm and blue for chromosome 16 q arm. Cell ID#20 of zygote z13 shows the loss of both chromosome 16 q arms while cell 21 demonstrates the addition of one chromosome 16 q arm. The blue box indicates the active chr16 centromere. Note the loss and gain of chromosomal material proximal to the cut site.

**Supplementary Figure 4.**
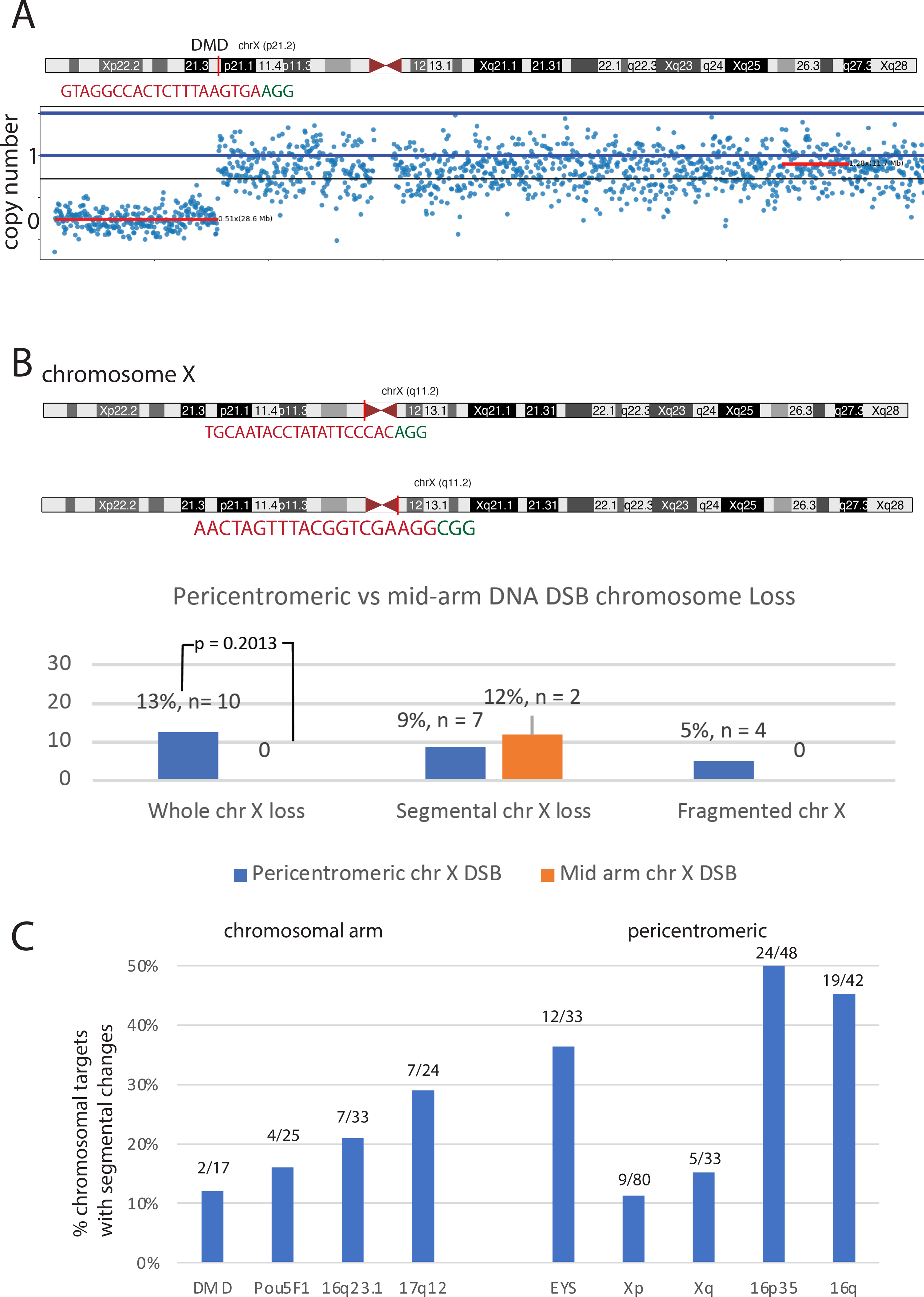
Mid-arm and pericentromeric DNA DSBs result in chromosomal changes. a. SNP analysis demonstrating segmental chromosome X loss at the targeted site on the mid-arm of chromosome X at the DMD locus. b. Location of pericentromeric gRNAs, and bar graph comparing chromosome loss after targeting the pericentromeric vs. mid-arm of chromosome X. c. Frequency of chromosomal changes per targeted chromosome. Numbers indicate the number of segmental changes/number of chromosomal targets.

**Table S1.** gRNA chromosome targets and sequences.

* Based on hg19 and hg38 annotation, this position is on the p arm of chromosome 16. Cleavage with this gRNA shows the functional centromere is proximal to this site.

** multiple secondary target sites with perfect match and a tolerated SNP on chr17:38487370, chr16:31991695, and chr16:32935225.

Bold: Pam site.

**Table S2**

Results of SNP Array Analysis and on-target Sanger sequencing analysis of Embryonic Cells after Cas9 Cleavage. Indels at the targeted genomic sequence by Cas9 gRNA based on Sanger sequencing. Samples with two complete homologous targeted chromosomes can have two different indels. Indels on maternal and paternal chromosome are separated by a dash (n/n) and indels where two chromosomes are present based on SNP arrays are indicated by asterisk (n*). These may be two homozygous indels or only one of the two is visible by PCR and Sanger sequencing. A size-neutral indel refers to a nucleotide insertion and deletion of equal value resulting in a net zero indel.

Detailed chromosomal content based on SNP array analysis.

**Table S3**

Cells or fragments that failed amplification or are not informative with regard to Cas9 activity because of chaotic aneuploidy or haploidy.

**Table S4**

Mapping of Cas9-induced break points as well as spontaneous chromosome breakage. Mapping was performed on samples with copy number transitions of 0-1 or greater, and 1-3 or greater. Samples with multiple gRNA targets on the same chromosome were not included in the calculation of DNA loss flanking the cut site.

**Table S5.** Primer sequences used for genotyping. Genomic coordinates according to hg38. Primers on 17q as well as on 16p arm can amplify from secondary sites as located in a repetitive region as the gRNA targeting these sites is.

## Notes

### Competing Interest Statement

J.X., D.M. and N.T. are employees and/or shareholders of Genomic Predictions Inc. D.E. and N.T. have applied for a patent on Cas9 mediated chromosome removal.

